# Oxytocin modulates social brain network correlations in resting and task state

**DOI:** 10.1101/2021.12.30.474596

**Authors:** Qingyuan Wu, Qi Huang, Chao Liu, Haiyan Wu

## Abstract

The effects of oxytocin (OT) on the social brain can be tracked upon assessing the neural activity in resting and task states, and developing a system-level framework for characterizing the state-based functional relationships of its distinct effect. Here, we contribute to this framework by examining how OT modulates social brain network correlations during resting and task states, using fMRI. First, we investigated network activation, followed by an analysis of the relationships between networks and individual differences. Subsequently, we evaluated the functional connectivity in both states. Finally, the relationship between networks across states was represented by the predictive power of networks in the resting state for task-evoked activities. The differences in the predicted accuracy between the subjects displayed individual variations in this relationship. Our results showed that the activity of the dorsal default mode network (DDMN) in the resting state had the largest predictive power for task-evoked activation of the precuneus network (PN) only in the OT group. The results also demonstrated that OT reduced the individual variation in PN in the prediction process. These findings suggest a distributed but modulatory effect of OT on the association between resting and task-dependent brain networks.

## 1. Introduction

Oxytocin (OT) has been commonly used to modulate human social behavior and neural activity of the social brain in both resting and task states (Brodmann et al., 2017; Horta et al., 2019; Hurlemann et al., 2010; Dirk Scheele et al., 2013; Wu, Liu, et al., 2020; Xin et al., 2021). The social brain comprises a set of regions, including the medial prefrontal cortex (MPFC), anterior cingulate cortex (ACC), temporal-parietal junction (TPJ), inferior frontal gyrus, and anterior insula (Adolphs, 2009; Barrett & Satpute, 2013; Frith, 2007; Stanley & Adolphs, 2013). Some task-dependent studies have revealed how OT impacts some of these brain regions. For example, the stress-induced response of the ACC is reduced by OT (Maier et al., 2019). Compared to the placebo (PL) group, the activation of the MPFC and ACC was also inhibited when patients with generalized social anxiety disorder saw an emotional face in the OT group (Freitas-Ferrari et al., 2010). About the resting state, several studies have found that OT changes the activation of networks consisting of regions that belong to the social brain (Jiang et al., 2021; Zheng et al., 2022; Zheng et al., 2021). Zheng et al. reported that OT leads to lower synchronization in the default mode network (DMN), which includes the posterior cingulate cortex (PCC), precuneus, MPFC, and TPJ (Zheng et al., 2022; Zheng et al., 2021). Another study found that the effective flow from the midline default network (including the PCC and precuneus) to the salience network (including the ACC and insula) was increased by OT (Jiang et al., 2021). Although some studies have focused on brain regions, the regions affected by OT belong to the default mode network or the salience network. Thus, these findings in the resting and task states suggest an effect of OT on these two networks.

Brain networks that support social functions may be comprised of individual differences related to distinct personality traits probed by scales, especially in different states (DeYoung, 2010; DeYoung et al., 2010; Laumann et al., 2015; Markett et al., 2018; Meyer et al., 2013). For example, in the resting state, individual functional networks may deviate from the group map and indicate variability across subjects (Kong et al., 2019; Laumann et al., 2015). Compared with the resting state, external stimuli in a task state modulate the connectivity of these brain networks (Cole et al., 2021). In recent years, researchers have gradually realized that the patterns of brain activity exhibited by individuals at rest could be a critical scaffold for people to perform behaviorally relevant tasks (Pang et al., 2022; Pezzulo et al., 2021; Reineberg et al., 2015; Shine et al., 2019; Tavor et al., 2016). As the general functional connectivity accounting for the resting-state and task-based fMRI drives reliable and heritable individual differences in functional brain networks (Elliott et al., 2019), exploring individual differences with a combination of brain networks across states and how those differences are affected, are emerging areas in neuroscience.

To the best of our knowledge, most previous studies have investigated the effects of OT on the resting- or task-state brain network separately at the group level. However, the question regarding the mechanism by which OT affects the scaffold effect of the spontaneous activity of the networks during the resting state remains unknown. A previous study has indicated that resting-state fMRI (rsfMRI) predicts individual differences in brain activity during task performance (Tavor et al., 2016). Subsequent studies have confirmed this prediction from rsfMRI to task-state fMRI (Bzdok et al., 2016; Cole et al., 2016; Mennes et al., 2010; Tavor et al., 2016). However, a previous study also suggested that brain network organization during a task is distinct from that in the resting state (Cole et al., 2014). Given individual variations in resting-state networks (RSNs), their influence on the brain network in subsequent tasks may be state-dependent. Although many studies have suggested that OT affects human social behavior and brain activity (Bartz et al., 2011; De Dreu et al., 2010; Hecht et al., 2017; Zhu et al., 2019), the lack of cross-state brain network investigation makes it impossible to reveal the mechanism of OT’s modulation effect of OT on the link between task-free and task-dependent brain networks.

However, recent studies have highlighted the subdivisions of classical brain networks. For instance, DMN have been divided into two sub-networks: the ventral default mode network (VDMN) and dorsal default mode network (DDMN) (Andrews-Hanna et al., 2010; Damoiseaux et al., 2008; Shirer et al., 2012). The VDMN is associated with memory-based reconstruction, whereas the DDMN is related to introspection on mental states and the evaluation of emotional valence (Chen et al., 2017; Lee et al., 2021). Additionally, the precuneus network (PN), including the precuneus, middle cingulate cortex, posterior inferior parietal lobule, and dorsal angular gyri, has been argued to be independent of the well-known DMN (Deng et al., 2019; Gilmore et al., 2015). Although the impact of OT on the DMN and the brain regions of the DMN has been discussed in many studies (Jiang et al., 2021; Zheng et al., 2022; Zheng et al., 2021), the specific modulatory effects of OT on connectivity among social sub-networks are largely unknown.

To investigate this, we tested the modulatory effects of OT on blood-oxygen-level-dependent (BOLD) signals during the resting state and during a task. Given that the impact of OT on the DMN has been confirmed in the resting and task states, and that the DMN is associated with self-related cues (as described above), we wished that the task included the perception of self-related cues. The face is an important social and emotional cue, and many studies have reported OT’s impact on the behavioral and neural activity when humans see faces (Lopatina et al., 2018; D. Scheele et al., 2013). Therefore, we designed a face perception task for the task state. During this period, we required participants to recognize whether a series of facial images were like themselves. Independent component analysis (ICA) was used to extract features in both states at the network level (Bzdok et al., 2016; Calhoun et al., 2008). After the networks were identified in both groups during task-dependent and resting states, we compared the correlations between the networks across the states. This allowed us to probe a critical question regarding the effect of OT on resting–task brain network correspondence. With the main aim of addressing whether OT could affect the relationship between networks in the resting and task states, we provide three sub-hypotheses in this work: 1) OT can decrease network connectivity in both states; 2) OT can influence resting–task brain network correspondence, particularly for brain regions related to social brain networks; and 3) OT may make human behavior patterns more consistent by reducing self-other differences in resting–task correspondence.

## 2. Method

### 2.1. Participants

Fifty-nine right-handed male participants (age range 20.9 ± 2.32 years) were recruited via an online recruiting system. All the participants had 13–18 years of education. Participants completed the screening form, and those who confirmed that they had no significant medical or psychiatric illness, were not using any medication, and were not drinking and/or smoking daily were included in the study. Smoking or drinking (except water) was prohibited for 2 h prior to the experiment. The participants provided written informed consent before each experiment and received a full debriefing on the completion of the experiment. The study was approved by the local ethics committee of the Beijing Normal University.

### 2.2. Resting and Task data sets

We used the resting and task state datasets collected from the same participants who were students at the universities of Beijing and who had been undergoing OT (OT group, including 30 participants) or PL (PL group, including 29 participants) manipulation. A single dose of 24 IU of oxytocin or placebo was administered intranasally. More details regarding drug administration have been described in one of our published studies (Wang et al., 2022; Wu, Feng, et al., 2020). The participants then underwent a resting-state scan (300s) before the task-state scan (240s).

For the resting dataset, we used fMRI data similar to the resting-state data that have been published for resting-state network studies only (Wu, Liu, et al., 2020; Zheng et al., 2022; Zheng et al., 2021). For the task dataset, we used the task fMRI data that were used for task fMRI analyses only (Wang et al., 2022), which provided the task descriptions. Specifically, we used a face perception task with morphed self and other faces (Wang et al., 2022). As the drug manipulation before scanning was a between-subjects design, and we collected the neural activation of each subject in both resting and task states, we had four states of fMRI dataset (OT-rest, OT-task, PL-rest, and PL-task).

Four types of face stimuli were generated: child self (morphing the participant’s face with one of two 1.5-year-old children’s faces with a neutral expression), child other (morphing the child’s face with one of two 23-year-old male faces), adult self, and adult other. Each participant completed two runs. Each run included three adult and three child blocks in a randomized sequence. Ten trials were performed for each block. The participants were asked to recognize whether the generated faces were similar to their own (Figure 1A).

**Figure 1:**
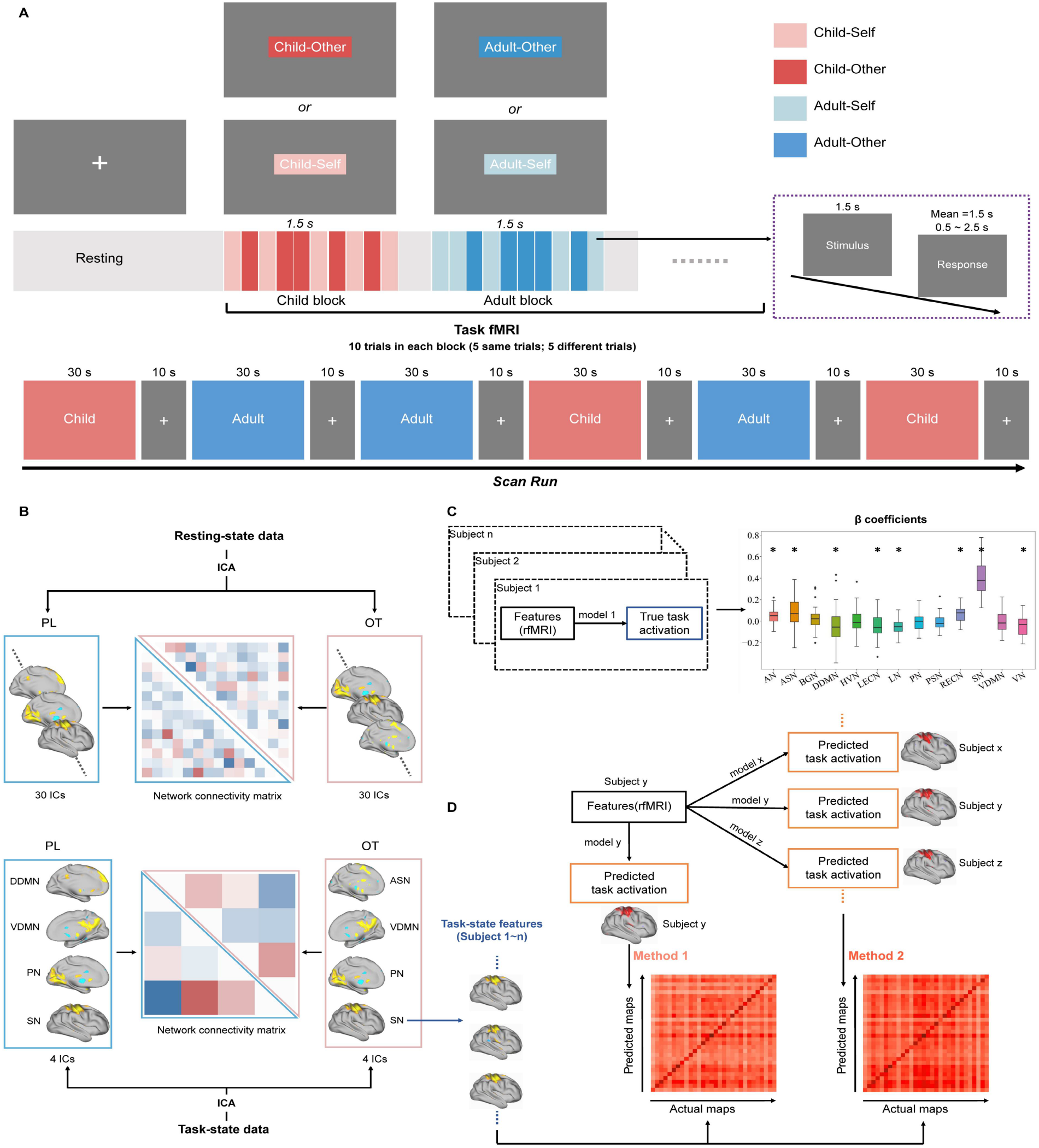
Diagrams representing our methods. A) Task procedure. Materials in the child block were morphed using a stranger child’s face with the participant’s (Self) face or another 23-year-old male’s (Other) face. Materials in the adult block were morphed using a stranger adult’s face and the participant’s (Self) face or 23-year-old male’s (Other) face. B) Feature extraction. We selected 30 features from the resting-state data and four features from the task state data from both groups. The correlation matrix was calculated for both states. C) Construction of the prediction models. In the training model step, we predicted features in task state by features in resting state and constructed a generalized linear model for each subject in this process. The outcomes were beta coefficients of each regressor (resting-state features). The beta values of features belonging to one network were averaged to be the network’s predictive power. D) The prediction model testing. We calculated the fitness between predicted task activation and the actual task activation for testing models. The first method (self-model method) to gain the prediction maps used one subject’s features (ICs extracted from rsfMRI data) and his own model. For this procedure, all participants shared one prediction map. The second method (other-model method) used one participant’s features (ICs extracted from rsfMRI data) and the model of others to generate a prediction map for each participant.

### 2.3. Materials

All participants had taken their frontal-face images with neutral expression three days before the scanning day. The morphed faces of the four experimental conditions were created based on these images and one of the two adult faces with a neutral expression (a 23-year-old male face) or one of the two 1.5-year-old child faces. We used Adobe Photoshop CS to standardize the photograph to black and white and excluded all features except the interior characteristics of the face. Then, we created stimuli with 50/50 morph of the two selected faces using the Abrosoft Fanta Morph (www.fantamorph.com) software [46, 47, 48]. Thirty calibration locations were used to transform the morphed face into a standard face space, and all the output morphed faces were resized to 300×300 dpi. All pictures were presented on a 17-inch Dell monitor with a screen resolution of 1024 × 768 pixels and 60 Hz refresh frequency. The visual angle of the face images was 4.3^◦^ ×4.6^◦^ and the mean luminance of the stimulus was 166 *cd/m*^2^.

### 2.4. MRI data acquisition

All images were acquired using a 3T Siemens Tim Trio scanner with a 12-channel head coil. Functional images employed a gradient-echo echo-planar imaging (EPI) sequence with following MRI scanner parameters: (40 *ms* TE, 2 *s* TR, 90^◦^ flip, 210 *mm* FOV, 3.5 mm*3.5 mm*4.2 mm voxel size, 128*128 matrix, 25 contiguous 5 *mm* slices parallel to the hippocampus and interleaved). We also acquired from all participants whole-brain T1-weighted anatomical reference images (2.15 *ms* TE, 1.9 *s* TR, 9^◦^ flip, 256 *mm* FOV, 176 sagittal slices, 1 *mm* slice thickness, perpendicular to the anterior-posterior commissure line).

### 2.5. fMRI data preprocessing

fMRI data preprocessing was performed using Statistical Parametric Mapping software (SPM12; Wellcome Trust Centre for Neuroimaging, London, UK). The functional image time series were preprocessed to compensate for slice-dependent time shifts, motion-corrected, and linearly detrended, then co-registered to the anatomical image, spatially normalized to the Montreal Neurological Institute (MNI) space, and spatially smoothed by convolution with an isotropic Gaussian kernel (FWHM=6 mm). fMRI data were band-pass filtered with a cutoff of 0.01–0.1 Hz. The white matter (WM) signal, cerebrospinal fluid (CSF) signal, and global signal, as well as the 6- dimensional head motion realignment parameters, the realignment parameters squared, their derivatives, and the squared of the derivatives were regressed.

### 2.6. Feature extraction

ICA was conducted using the Group ICA fMRI Toolbox (GIFT) to extract the brain activation features (Rachakonda et al., 2007). For rsfMRI data, in the first step, GIFT estimated the number of independent components (ICs) to be extracted from the preprocessed rsfMRI data. Second, we calculated the correlation coefficients between the ICs and RSN templates (Reineberg et al., 2015). These templates included 14 brain networks (SI Figure 4): the basal ganglia network (BGN), visuospatial network (VN), right executive control network (RECN), VDMN, DDMN, sensorimotor network (SN), anterior salience network (ASN), auditory network (AN), higher visual network (HVN), left executive control network (LECN), language network (LN), PN, posterior salience network (PSN), and primary visual network (PVN) (Shirer et al., 2012). Each IC was compared with all brain network templates and labelled based on the most similar network defined in the template. Finally, 30 ICs with correlation coefficients greater than 0.1 were separately retained as predictors for both groups (Figure 1B) (Reineberg et al., 2015).

For the task-state fMRI data, we estimated and analyzed the data using the method described in the previous paragraph. Next, to confirm that the activation of ICs was aroused by our task, we compared the time course of ICs and task GLM activation maps. Specifically, the correlation analysis was performed between the time courses of these ICs with the modelled time course (the beta maps) for all groups in the task of the 1st level SPM.mat The selected regressors were “self-child*bf(1)”, “other-child*bf(1)”, “self-adult*bf(1)”, and “other-adult*bf(1)” time courses (Rachakonda et al., 2007). ICs that were most closely related to task activation (correlation coefficient *>* 0.7) were chosen as features. Finally, four ICs were chosen as features for both the PL and OT groups (Figure 1B). We labeled them using a template.

### 2.7. Predictive Model

To predict task-state fMRI by rsfMRI, following the group ICA described above, we used dual regression to reconstruct the subject-specific spatial z-score maps and subject-specific time course of each IC during both states (Beckmann et al., 2009). Specifically, subject-specific time series were generated by regressing the group-averaged set of spatial maps on the subject’s 4D spatiotemporal dataset. Then, the subject-specific components (whole-brain images) were obtained by regressing those subject-specific time series into the same 4D dataset.

For each subject, all ICs extracted from the rsfMRI were applied to predict each IC in the task-state fMRI data. Therefore, we constructed four multiple linear regression models for each subject; each model included 30 regression coefficients (Tavor et al., 2016). The coefficients of ICs belonging to the same network template were averaged to intuitively observe the predictive power of each RSN (Figure 1C). We then used a one-sample t-test to assess whether RSNs activation could significantly predict task-state brain network activation.

We also examined the predictive effect of individual differences in psychometric scale scores on the activation of RSNs. Previous studies have indicated that the effect of OT on brain activity is modulated by emotional states and individual variability(Alcorn III et al., 2015; Hecht et al., 2017; Xin et al., 2020). Thus, we measured whether OT could affect the associations between brain network activation and subjective emotional states as well as personality traits. Emotional states were measured using the Positive and Negative Affect Schedule (PANAS) (Thompson, 2007), and the Big Five scale (Bozionelos, 2004) was used to detect participants’ personality traits. Regression analysis was performed using generalized estimated equations to predict the number of ICs in the specific networks associated with social brain networks, including the BGN, RECN, VDMN, DDMN, SN, ASN, LECN, and PN. The predictors in the generalized estimated equations were the PANAS and Big Five scores measured after OT and PL manipulation (Gaviria et al., 2021).

### 2.8. Functional connectivity analysis

We also analyzed the relationship between networks in both resting and task states. The ICs sharing the same label were averaged at the voxel level to indicate brain network activation. We then calculated the Pearson coefficient matrices of all the networks for both groups in the two states. Because the two groups of extracted ICs differed in the task state, an independent t-test was only performed to probe the influence of OT on the network correlation at rest. P-values were adjusted using Benjamini and Hochberg’s false discovery rate correction for multiple comparisons (Gaviria et al., 2021).

### 2.9. Model test

The coefficient of determination was regarded as an indicator for estimating the predictive power of our models and was compared with that of the null model (all regressors were equal to 1). We then performed a one-sample t-test for the indicators of each model and assessed their performance (Chatterjee & Hadi, 1986).

One way to assess the model performance was to obtain the prediction accuracy by comparing the actual task maps with the prediction activation maps. Prediction activation maps can be derived using these two methods. The first method combined the rsfMRI data of subject X with his own model (Self-model, SM) and compared this prediction map with the subjects’ actual task maps (Tavor et al., 2016). The second method combined the rsfMRI data of subject X with the model of subject Y (Other-model, OM) to obtain a prediction map for subject Y. In other words, each subject had a specific prediction map using this method. The correlation coefficients between the prediction maps and actual task-state activity maps were considered as the predicted accuracies (Figure 1D). Each subject’s RSNs activation was used to predict task-related brain activation in all subjects. A Pearson’s correlation matrix was obtained for each IC (Figure 1D). The diagonal elements indicate self-predicted accuracy (X’s resting brain activation predicts X’s task brain activation), and the extra-diagonal elements indicate other-predicted accuracy (X’s resting brain activation predicts Y’s task brain activation). Thus, the difference between the two accuracies represents the individual variance (self-other difference). The percentage of the self-other difference relative to the average of other-predicted accuracy could indicate the relative size of the difference (Tavor et al., 2016). We then compared the percentage of self-other differences between the PL and OT groups using an independent sample t-test.

## 3. Results

### 3.1. Behavioral results

We tested two indices that indicated the subjects’ behavioral performance: 1) the accuracy of recognizing whether a morphed face was similar to the subject himself, and 2) the reaction time for subjects to make judgments. We calculated the two indices for subjects in four conditions (OT-child, OT-adult, PL-child, and PL-adult) separately. Two indices for each subject were separately put into a 2-way mixed analysis of variance (ANOVA) using treatment groups (OT vs. PL) as between-subject factors and facial conditions (self vs. other, and child vs. adult face) as within-subject factors.

For accuracy, there was only a significant main effect of facial conditions ((*F* (1, 57) = 54.6716, *p* < 0.001, *η*_p_^2^ = 0.4949)), but no significant effect was observed on treatment (OT and PL groups) and interaction (*p* >0.05). For RTs, we did not find any significant effects of the treatment. These results indicated that OT did not influence behavioral performance in the face perception task. Further details are provided in one of our published studies (Wang et al., 2022).

### 3.2. The effect of OT on the resting brain networks is modulated by individual differences

We first examined the effects of OT on brain network activity at rest. Networks correlated with the emotional or social brain (ASN, BGN, DDMN, LECN, RECN, SN, PN, and VDMN) were selected for the following analysis. To test whether the emotional and personal traits were correlated with brain activity, we measured the scale scores after drug administration. A significant effect of OT on the presentation of emotion and personality in brain networks may indicate that the subjective affective rating scores (measured by PANAS) and personality scores (measured by Big-Five) could predict the occurrence of brain networks. The results showed that approximately all emotional scores were associated with bilateral executive control network activity, but only the extraversion scores on the personality scale could predict BGN and LECN activity in the OT group. However, for the PL group, personality scores had significant predictive effects on multiple brain networks, while emotional scores were almost completely unrelated to the activity of brain networks (Figure 2).

**Figure 2:**
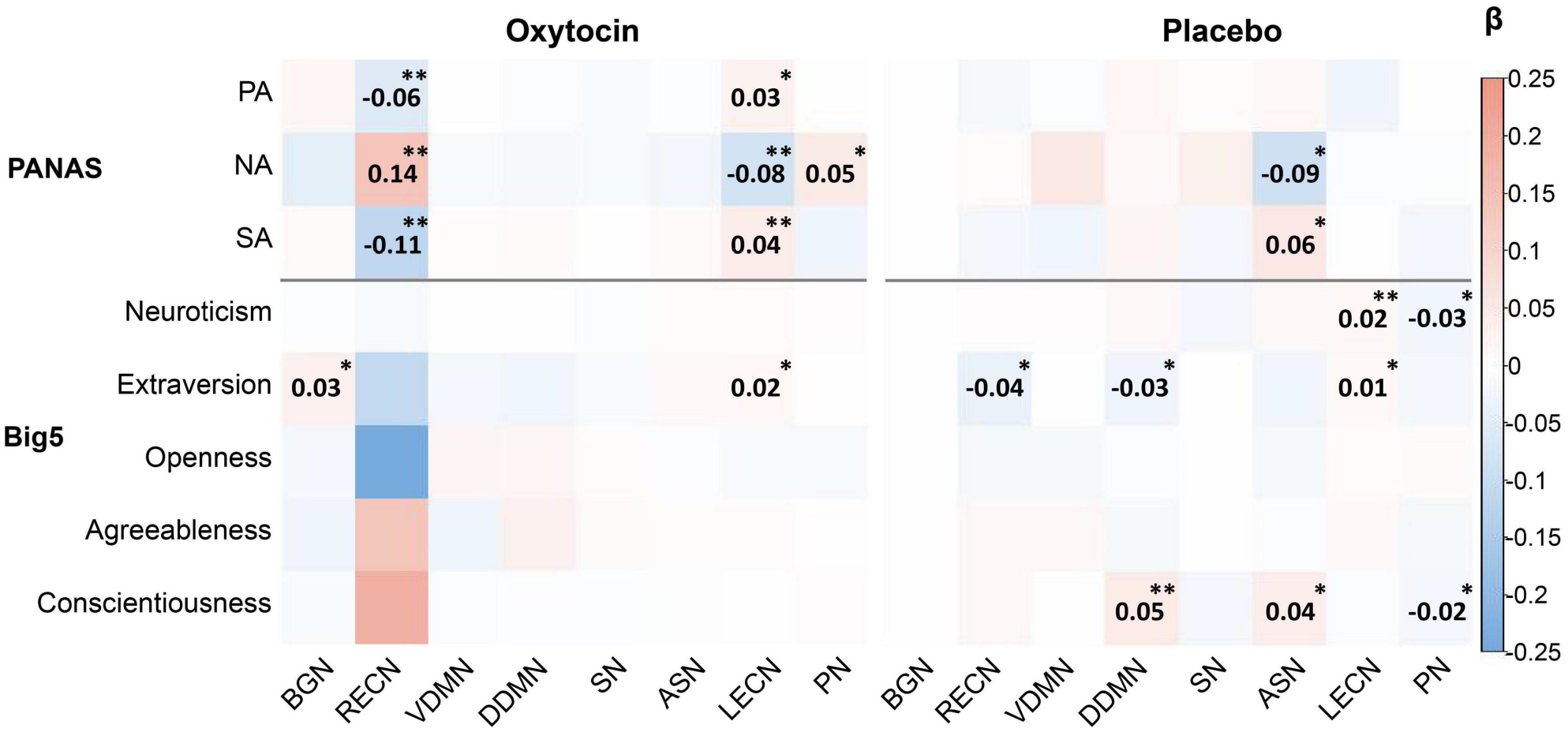
PANAS and Big-Five scores’ associations with brain networks across the two groups. Beta values of generalized estimated equations-based associations between the occurrence of the ICs of interested network and scale scores, including affective scores (positive (PA) and negative (NA)) and personality scores. ^∗^*p* < 0.05, ^∗∗^*p* < 0.01.

### 3.3. Network correlation in resting and task state

Next, we explored the influence of OT on the relationship between the brain networks by calculating the functional connection matrix in the two states respectively. The results showed that OT not only affected the connection strength but also the direction of correlation between brain networks in the resting state (see SI Figure 1 and Table 1). To better visualize the effects of OT, we compared the matrix of the OT group with that of the PL group (Figure 3A and Table 1).

**Figure 3:**
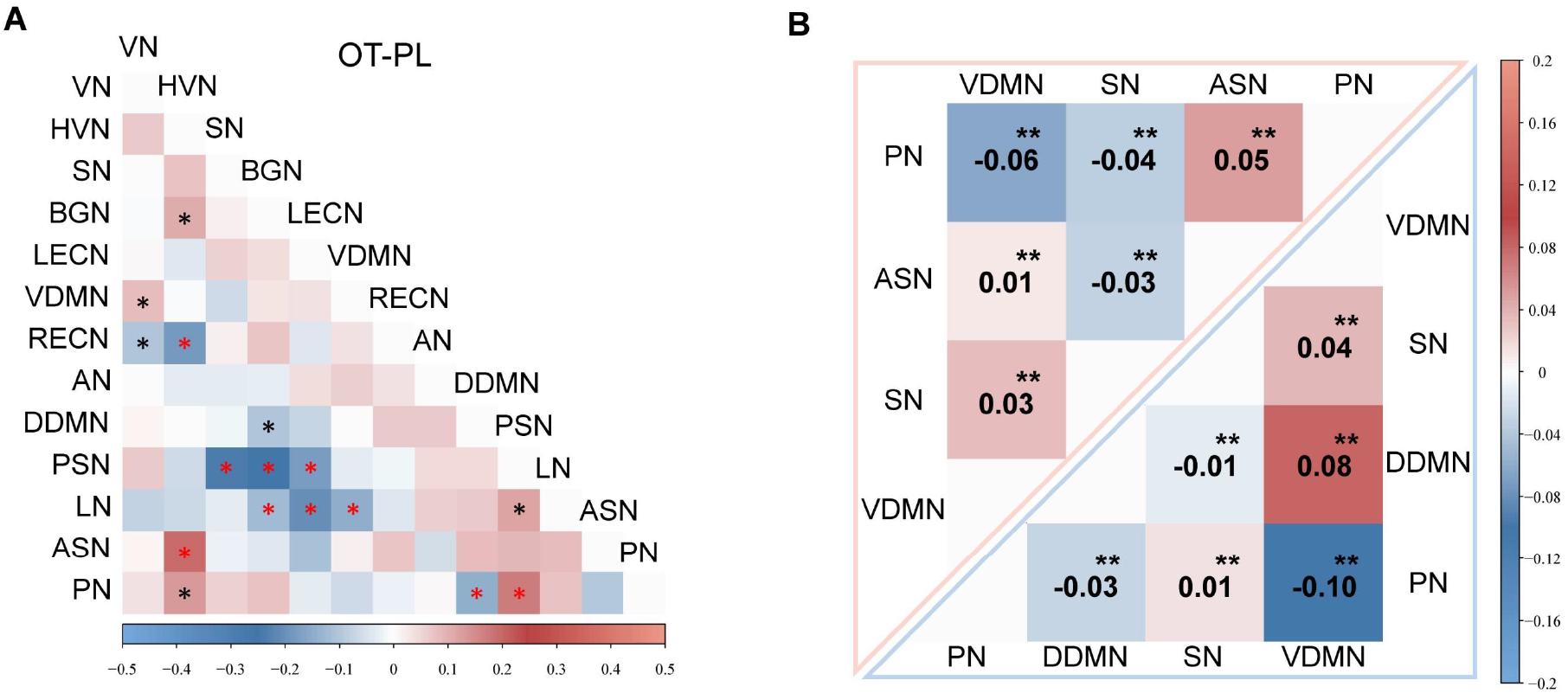
The difference of correlation matrix between brain networks (OT-PL). A) The difference between two group-level matrixes. The red star indicates *p* < 0.001, and the black star indicates *p* < 0.05. B) The task state brain network correlation matrix. Group-level correlation matrix of features extracted from the task fMRI data. *pFDR* adjustment for multiple comparisons. ^∗∗^*p* < 0.01

**Table 1:** RSNs correlation coefficients

|  | **OT** | **PL** | **t-score** | **$p_{FDR}$** |
| --- | --- | --- | --- | --- |
| PN-DDMN | -0.12 | 0.02 | -5.08 | 0.000 |
| HVN-ASN | 0.08 | -0.12 | 5.29 | 0.000 |
| PSN-SN | -0.05 | 0.19 | -7.16 | 0.000 |
| PSN-BGN | -0.15 | 0.10 | -7.00 | 0.000 |
| PSN-LECN | -0.14 | 0.04 | -4.55 | 0.000 |
| LN-LECN | -0.27 | -0.06 | -5.05 | 0.000 |
| LN-VDMN | -0.06 | 0.09 | -5.03 | 0.000 |
| HVN-RECN | 0.01 | 0.19 | -4.92 | 0.000 |
| PSN-PN | 0.09 | -0.09 | 5.17 | 0.000 |
| DDMN-BGN | -0.19 | -0.09 | -3.36 | 0.002 |
| LN-BGN | -0.12 | 0.01 | -3.58 | 0.002 |
| VDMN-VN | 0.01 | -0.07 | 3.22 | 0.002 |
| PSN-LN | 0.11 | 0.00 | 3.25 | 0.002 |
| PN-HVN | -0.09 | -0.23 | 3.19 | 0.002 |
| HVN-BGN | -0.01 | -0.12 | 3.16 | 0.003 |
| RECN-VN | -0.08 | 0.02 | -2.98 | 0.004 |

*Note:* Means, and two-tailed t-test (direction: OT - PL) results of the comparisons on functional connectivity in the resting state.

For the task-state dataset, the ICs correlated with the task were labeled as PN, SN, ASN, and VDMN in the OT group, and those for the other group were labeled as PN, SN, DDMN, and VDMN. The results showed a difference in the activity of DDMN between the groups, but on difference for VDMN. This finding was consistent with the resting-state results, which further indicated that the VDMN and DDMN may be affected differently by OT. Furthermore, we explored task-state brain network correspondence. The results indicated that the correlation between PN and ASN was opposite to that between PN and the DDMN (Figure 3B). This is distinct from the pattern observed during the resting state. In this state, the correlation between PN and ASN was the same as the correlation between PN and DDMN in both groups (SI Figure 1).

### 3.4. Associations between resting-state and task-state networks

Figure 4 shows the contribution of resting ICs to recovering the activation of task ICs. All brain networks in the task state, except for PN in the OT group, could be best predicted themselves in the resting state. In the PL group, the strongest predictor of PN in the state. However, under the influence of OT, the predictive power of the DDMN was larger than that of the PN itself (*betaDDMN* = 0.56, *betaPN* = 0.26, *t* (118) = 3.00, *p* = 0.003, box plots in Figure 4). This finding suggested that OT might strengthen the scaffolding effect of DDMNs on PN. Generally, there was less correlation between the RSNs and the task-state network in the OT group.

**Figure 4:**
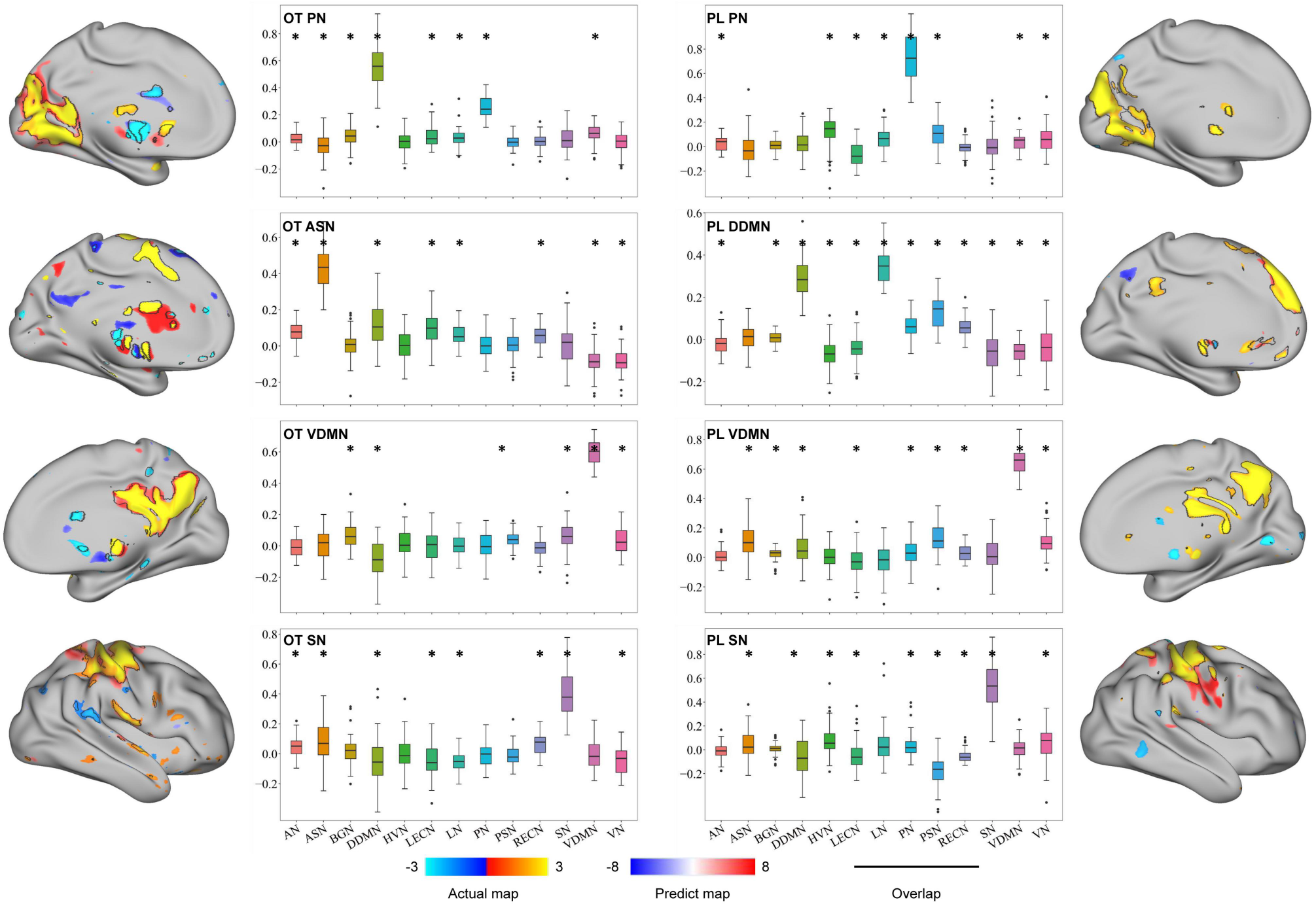
Predicting individual task activation maps. Beta coefficients (vertical coordinates) averaged across ICs that belong to one brain network indicate the contribution of the resting-state networks in predicting task-state features. For the most of features, the network’s self-predictive power was the strongest, except for the PN under the OT group. The brain plots show actual and predicted task contrast maps for all features that are close to our task in two subjects (one for the OT group and one for the PL group). ^∗^*pFDR* < 0.05.

We used the ICs from the rsfMRI data to forecast the four task-state ICs and obtained four prediction maps from each participant after model construction. We measured the recovery performance by comparing the predicted maps with actual maps. The actual and predicted maps of the main regions of each IC showed a significant rate of overlap (Figure 4). Our approach was also able to detect inter-subject variability in both groups. The model correctly recovered activation in areas that had no expression, on average(Figure 5). As a result, we investigated individual variations in network interactions across resting and task states. The subsection, ”Model performance and inter-subject variation” provides more detail about the predictive accuracy and individual variation.

**Figure 5:**
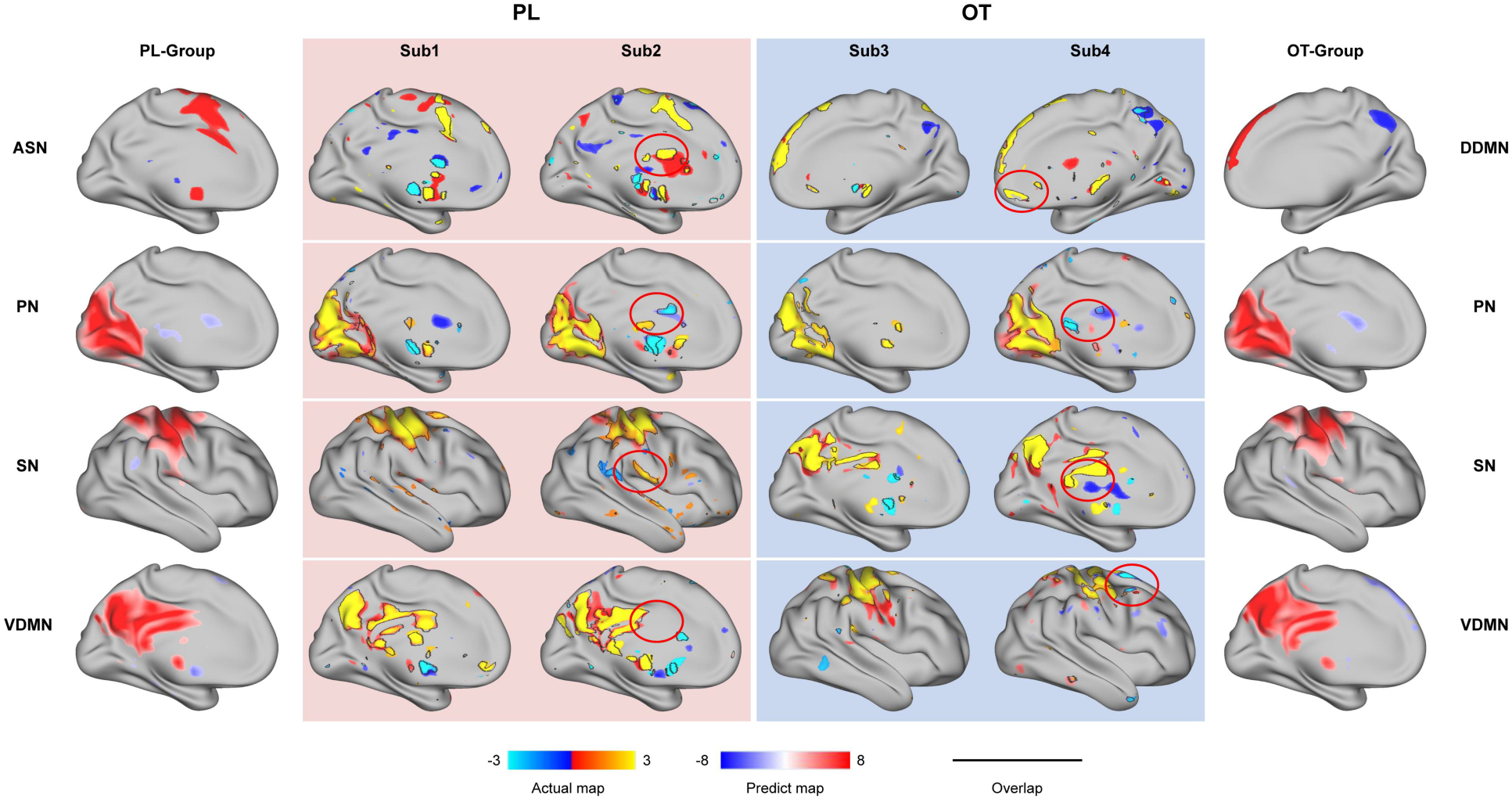
Personalized predictive maps. In both groups, our model showed the ability to detect inter-subject variability. The brain maps with white context indicate the group-averaged activation maps of networks, and the other four columns with color context show the actual and predicted activations of two subjects in one group (OT or PL). Although the main region was similar to the group activation, the model captured the specific clusters that were not showed in the group.

### 3.5. Model performance and the inter-subject variation

Two indices were used to test the model: The first was the coefficient of determination *R*^2^, which represents how many variations could be explained by the models. Although the networks for prediction (resting-state ICs) and the modelling process were the same, we found that the variation in different task ICs explained by resting ICs varied greatly. Interestingly, the models performed best when interpreting DDMN(*R*^2^ = 0.39) but worst when interpreting ASN(*R*^2^ = 0.23). In addition, no significant differences in the interpretability of the same brain network were found between groups (SI Figure 2, Table 2).

**Table 2:** Coefficients of determination.

|  | **Mean R square std** |  |
| --- | --- | --- |
| PL DDMN | 0.39 | 0.09 |
| PL PN | 0.31 | 0.10 |
| PL VDMN | 0.32 | 0.08 |
| PL SN | 0.24 | 0.10 |
| OT ASN | 0.23 | 0.07 |
| OT PN | 0.33 | 0.10 |
| OT VDMN | 0.33 | 0.07 |
| OT SN | 0.23 | 0.07 |

*Note:* Means and standard deviations results of the coefficients of determination.

Another index is prediction accuracy (see ”Model test”). We examined whether the two methods (see the Model Test) yielded different results. In most situations, the self-model method performed better than the other model methods, except for the VDMN in the PL group and ASN in the OT group (Figure 6A, SI Table 1). Additionally, only PN showed significantly higher accuracy in the OT group than in the PL group for both methods (Figure 6A, SM: *t* (58) = 2.79, *p* = 0.007; OM: *t* (58) = 2.96, *p* = 0.004).

**Figure 6:**
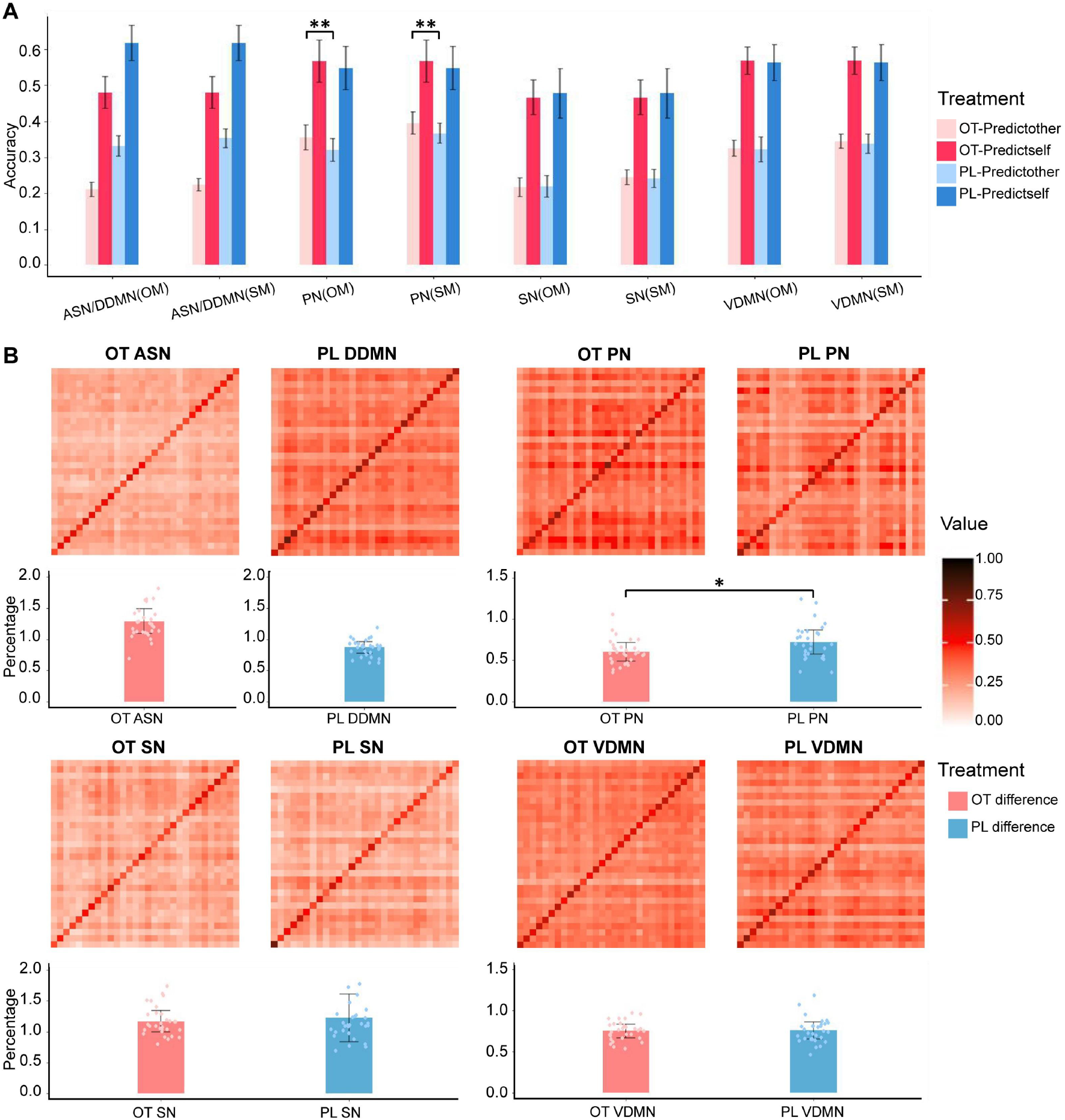
Performance of the model and specificity of the individual predictions. A) This figure shows the accuracy of using one’s own model (SM) and others’ models (OM) to predict one’s own (deep color) and others’ task-evoked network activation (light color) for both conditions. B) Pearson correlation matrix between actual (columns) and predicted activations (rows). The elements on the diagonal represent the accuracy of self-prediction, while those extra-diagonal elements show the other-prediction accuracy. The bar plot, which was produced by the method using other-models, indicates the difference between the accuracy of using participants’ resting-state data to predict their task activation and predicting others’ activation (as a percentage relative to the average accuracy of predicting others).

Additionally, compared to the PL group, there was a smaller difference in the accuracy between predicting the own-task and other-task activation maps in the OT group. Both methods showed that only PN displayed a significantly higher accuracy in predicting others’ task maps in the OT group than in the PL group (Table 3, Figure 6A). The difference in the accuracy of PN in the OT group was lower than that in the PL group when performing the other models (*t* (58) = -2.55, *p* = 0.014, Figure 6B). Meanwhile, the SN and VDMN also showed this trend, although the difference was not significant. These results suggest that OT may reduce interindividual differences in PN. To further confirm this finding, we extracted ICs from the data of all subjects (including the OT and PL groups) and repeated the analysis that produced the results shown in Figure 6, and we obtained the same results as Figure 6 (SI Figure S3). However, this effect was not observed when self-modeling was used.

**Table 3:** Predictive accuracy of others

|  | OT | PL | t-value | p |
| --- | --- | --- | --- | --- |
| PN(Self-model) | 0.37 | 0.40 | -2.79 | 0.007 |
| PN(Other-model) | 0.32 | 0.36 | -2.96 | 0.04 |
| VDMN(Self-model) | 0.34 | 0.34 | -0.83 | 0.408 |
| VDMN(Other-model) | 0.32 | 0.33 | -0.28 | 0.783 |
| SN (Self-model) | 0.22 | 0.22 | 0.21 | 0.700 |
| SN (Other-model) | 0.22 | 0.21 | 0.42 | 0.834 |

*Note:* Means, and two-tailed t-test (direction: OT - PL) results of the comparisons on predictive accuracy.

## 4. Discussion

To the best of our knowledge, although the effect of OT on neural activity in both resting and task states has been well discussed, this work is the first to investigate how OT affects brain network activation and the relationships of these networks across resting and task states. In addition to confirming the effect of OT on brain networks during both resting state and face perception tasks; we further assessed its impact on the scaffolding mechanism of RSNs for task-evoked network activation. Our study provided four promising preliminary results. First, compared with the PL group, there was decreased functional connectivity among the networks in the OT group. Second, in the OT group, the resting-state activity of the DDMN showed the largest predictive power for the task-evoked brain activity of the PN, but not for the PL group. Finally, OT reduced self-other differences in PN when predicting task activation. Overall, our findings provide new evidence for dynamic and interactive modulation of brain-wide networks by OT.

### 4.1. DDMN has the highest predictive power for task-evoked activity of PN for the OT group

Studies have shown a close relationship between PN and DMN, but with mixed results regarding whether PN is a part of the DMN (Fransson & Marrelec, 2008; Utevsky et al., 2014). Indeed, some studies have shown that the precuneus is the functional core of the DMN(Deng et al., 2019; Gilmore et al., 2015). However, recent studies do not support this idea and indicate that the PN is distinct from the DMN (Gilmore et al., 2015). A meta-analysis of task-based fMRI and resting-state functional connectivity studies identified the PN (also called ‘parietal memory network’) as broadly involved in the formation of memory and personal experience of novelty (Gilmore et al., 2015). Another fMRI study demonstrated the distinct functional roles of the PN and DMN in processing context-rich information (Deng et al., 2019). Our findings may add another piece of evidence on the overlap or interaction between PN and DMN.

In our results, we found that PN was negatively correlated with DDMN and VDMN during the task, regardless of whether it was affected by OT. The functional connectivity between the PN and DDMN at rest was reversed in the OT group compared to that in the PL group (Figure 3A, SI. Figure 1). Additionally, DDMN at rest had a stronger predictive power for PN in the task state than resting PN itself in the OT group, but no significant predictive power was observed in the PL group (Figure 4). This was the only case in which the network’s performance at rest was not the best predictor of its activation during the task. Together, these results suggest that dynamic network changes depend on drug administration; for example, OT changes the way in which the DDMN works in the task. In the PL group, the DDMN played a direct role in the task. In the OT group, the DDMN participated in the task by modulating the activation of the PN, rather than acting directly. Additionally, combining our current work with previous studies (Wang et al., 2022; Wu, Feng, et al., 2020), which showed that OT influenced neural activation in both states, we consider both changes in resting and task states lead to a change in the correlation between the resting-state and task-related networks.

### 4.2. OT reduced individual difference in PN prediction

Our study provides two results regarding the impact of OT on individual variations in PN activation in both the resting and task states. First, OT may reduce the predictability of PN activation when using the Big-Five scale scores to predict the activity of PN (Figure 2). The second is reflected in the process of using rsfMRI data to predict network expression in a task state. The result indicates that the self-other difference (i.e., the distinction between using subject X’s resting-state data to predict X’s task data and to predict subject Y’s task data) of PN was significantly decreased by OT (Figure 6B). These findings are likely opposite to previous studies, which suggest that the impact of OT on individuals displays vast individual variability (Alcorn III et al., 2015; Hecht et al., 2017; Koch et al., 2016; Kumsta & Heinrichs, 2013). For instance, some studies have reported that brain activation was significantly correlated with personal characteristics in the OT group, while the correlation was not significant in the PL group (Alcorn III et al., 2015; Hecht et al., 2017; Koch et al., 2016). We propose that the reduced individual variation makes the correlations between personal characteristics and brain activity easier to observe. Notably, our resting/task-state prediction of the two groups focused more on task-brain prediction than on individual differences per se. This may provide new insights for understanding the mechanism by which OT affects human brain activity during tasks, depending on brain activity during the resting state.

In the OT group, we found a smaller self-other difference when predicting the task network activation, which was only significant for PN. This may be due to the specific role of PN in our self-other face perception task. Cavanna and Trimble (2006) described a series of studies in which the precuneus was activated more for self-relevant than self-irrelevant personal traits (Kircher et al., 2000; Kjaer et al., 2002; Lou et al., 2004), and more for a first-person than a third-person perspective (Den Ouden et al., 2005; Vogeley et al., 2001; Vogeley & Fink, 2003). Therefore, when subjects recognize whether a face stimulus is similar to the face of the subjects themselves, OT may have the greatest impact on the most important brain network, PN. Thus, this provides further evidence that task-evoked and connectivity data are essential for studying the resting– task predictable networks.

## 5. Limitation and future direction

The present study provides the first evidence that OT modulates the resting–task brain network in a self-other face perception task. It remains unknown whether our results can be generalized to other social tasks. In addition, the reason for different networks displaying distinct predictability in our study remains unknown. Additionally, the sample size and the fact that our sample consisted of only men may also limit the generalizability of our results. For example, we cannot investigate all profiles of OT effects on behavioral and brain networks that can be predicted by resting state, with one representative task. Perhaps, different types of tasks (e.g., social and cognitive tasks) may result in specific OT effects on resting–task brain prediction. Future studies are required to test the correlations between resting and task-related brain networks after OT administration in multiple-task settings.

Another topic worth exploring is the consistency between people’s subjective perceptions and their brain activity in different states. Accumulating evidence suggests that OT increases prosocial behavior (Bartz et al., 2011; Carter, 2014; De Dreu & Kret, 2016), and our findings indicate that individual differences in networks decrease in the OT group. Previous studies on brain–heart interactions have reported that information flows more from the heart to the brain during a relaxed state, such as sleep (Min & Wang, 2017), but more information flows from the brain to the heart as the consciousness arousal level increases (Abukonna et al., 2013). Other studies have shown higher brain activation synchronism when people better understand others (Simony et al., 2016; Zadbood et al., 2017). The brain activity patterns of friends are more similar to those of strangers (Hyon et al., 2020; Parkinson et al., 2018). Since all these findings demonstrated individual insights into their neural system, does this consistency generalize to the prediction process? For example, whether the accuracies of individuals in the resting state in predicting their performance in a task are consistent with the predictability of their task brain activity using their rsfMRI or not. Which brain regions or networks display more consistency with subjective human perceptions? What factors could affect this consistency? We believe that our study may inspire future research regarding factors such as cue presentation sequence, mood, and social information, contributing to the enhancement of resting–task network prediction.

## 6. Conclusion

In this study, we collected fMRI data at rest and task states in both OT and PL groups to examine how OT impacts 1) the co-activation of brain networks in two states, 2) the predictive power of RSNs for task-evoked network activity, and 3) the individual variation reflected in the predictive process. As a result, we found that the dorsal default mode network (DDMN) was decreased by OT in the resting state; OT changed the greatest predictor for the task-evoked activation of the precuneus network (PN) from PN to DDMN (in the resting state); OT reduced the individual variation of PN, specifically, the difference in accuracy between predicting subjects’ own and others’ PN task activation. We believe that these findings together reflect OT’s distinct influence on diverse brain networks across different states and their relationships. The current study elucidated the mechanism by which OT affects the scaffolding mechanism of networks in the resting state during task activation.

## Abbreviations

OT: oxytocin
PL: placebo
fMRI: functional magnetic resonance imaging
rsfMRI: resting-state fMRI
BOLD: blood-oxygen-level-dependent
MPFC: medial prefrontal cortex
ACC: anterior cingulate cortex
TPJ: temporal-parietal junction
PCC: posterior cingulate cortex
ICA: independent component analysis
ICs: independent components
RSNs: resting-state networks
DMN: default mode network
VDMN: ventral default mode network
DDMN: dorsal default mode network
PN: precuneus network
BGN: basal ganglia network
LECN: left executive control network
RECN: right executive control network
PN: precuneus network
SN: sensorimotor network
ASN: anterior salience network
AN: auditory network
HVN: higher visual network
LN: language network
PSN: posterior salience network
PVN: primary visual network
PANAS: positive and negative affect schedule

## Acknowledgements

This work is funded by Natural Science Foundation of Guangdong Province (2021A1515012509), SRG of University of Macau (SRG2020-00027-ICI), Shenzhen-Hong Kong-Macao Science and Technology Innovation Project (SGDX 2020110309280100), Science and Technology Development Fund (FDCT) of Macau (0127/2020/A3), National Natural Science Foundation of China (NSFC) (U1736125), the Major Project of National Social Science Foundation (19ZDA363), the Beijing Municipal Science and Technology Commission (Z151100003915122), and Open Research Fund of the State Key Laboratory of Cognitive Neuroscience and Learning (CNLYB2002).

## References

Abukonna, A., Yu, X., Zhang, C., et al. (2013). Volitional control of the heart rate. Int J Psychophysiol, 90(2), 143–148. https://doi.org/10.1016/j.ijpsycho.2013.06.021

Adolphs, R. (2009). The social brain: neural basis of social knowledge. Annu Rev Psychol, 60, 693–716. https://doi.org/10.1146/annurev.psych.60.110707.163514

Alcorn III, J. L., Green, C. E., Schmitz, J., et al. (2015). Effects of oxytocin on aggressive responding in healthy adult males. Behavioural pharmacology, 26(8 0 0), 798.

Andrews-Hanna, J. R., Reidler, J. S., Sepulcre, J., et al. (2010). Functional-anatomic fractionation of the brain’s default network. Neuron, 65(4), 550–562. https://doi.org/10.1016/j.neuron.2010.02.005

Barrett, L. F., & Satpute, A. B. (2013). Large-scale brain networks in affective and social neuroscience: towards an integrative functional architecture of the brain. Curr Opin Neurobiol, 23(3), 361–372. https://doi.org/10.1016/j.conb.2012.12.012

Bartz, J. A., Zaki, J., Bolger, N., et al. (2011). Social effects of oxytocin in humans: context and person matter. Trends Cogn Sci, 15(7), 301–309. https://doi.org/10.1016/j.tics.2011.05.002

Beckmann, C. F., Mackay, C. E., Filippini, N., et al. (2009). Group comparison of resting-state FMRI data using multi-subject ICA and dual regression. Neuroimage, 47(Suppl 1), S148.

Bozionelos, N. (2004). The big five of personality and work involvement. Journal of Managerial Psychology.

Brodmann, K., Gruber, O., & Goya-Maldonado, R. (2017). Intranasal Oxytocin Selectively Modulates Large-Scale Brain Networks in Humans. Brain Connect, 7(7), 454–463. https://doi.org/10.1089/brain.2017.0528

Bzdok, D., Varoquaux, G., Grisel, O., et al. (2016). Formal Models of the Network Co-occurrence Underlying Mental Operations. PLoS Comput Biol, 12(6), e1004994. https://doi.org/10.1371/journal.pcbi.1004994

Calhoun, V. D., Kiehl, K. A., & Pearlson, G. D. (2008). Modulation of temporally coherent brain networks estimated using ICA at rest and during cognitive tasks. Human Brain Mapping, 29(7), 828–838. https://doi.org/10.1002/hbm.20581

Carter, C. S. (2014). Oxytocin pathways and the evolution of human behavior. Annu Rev Psychol, 65, 17–39. https://doi.org/10.1146/annurev-psych-010213-115110

Chatterjee, S., & Hadi, A. S. (1986). Influential observations, high leverage points, and outliers in linear regression. Statistical science, 379–393.

Chen, J. Y. E., Glover, G. H., Greicius, M. D., et al. (2017). Dissociated Patterns of Anti-Correlations with Dorsal and Ventral Default-Mode Networks at Rest. Human Brain Mapping, 38(5), 2454–2465. https://doi.org/10.1002/hbm.23532

Cole, M. W., Bassett, D. S., Power, J. D., et al. (2014). Intrinsic and task-evoked network architectures of the human brain. Neuron, 83(1), 238–251. https://doi.org/10.1016/j.neuron.2014.05.014

Cole, M. W., Ito, T., Bassett, D. S., et al. (2016). Activity flow over resting-state networks shapes cognitive task activations. Nat Neurosci, 19(12), 1718–1726. https://doi.org/10.1038/nn.4406

Cole, M. W., Ito, T., Cocuzza, C., et al. (2021). The Functional Relevance of Task-State Functional Connectivity. J Neurosci, 41(12), 2684–2702. https://doi.org/10.1523/JNEUROSCI.1713-20.2021

Damoiseaux, J. S., Beckmann, C. F., Arigita, E. J. S., et al. (2008). Reduced resting-state brain activity in the “default network” in normal aging. Cerebral Cortex, 18(8), 1856–1864. https://doi.org/10.1093/cercor/bhm207

De Dreu, C. K., Greer, L. L., Handgraaf, M. J., et al. (2010). The neuropeptide oxytocin regulates parochial altruism in intergroup conflict among humans. Science, 328(5984), 1408–1411.

De Dreu, C. K., & Kret, M. E. (2016). Oxytocin conditions intergroup relations through upregulated in-group empathy, cooperation, conformity, and defense. Biological psychiatry, 79(3), 165–173.

Den Ouden, H. E., Frith, U., Frith, C., et al. (2005). Thinking about intentions. Neuroimage, 28(4), 787–796.

Deng, Z., Wu, J., Gao, J., et al. (2019). Segregated precuneus network and default mode network in naturalistic imaging. Brain Struct Funct, 224(9), 3133–3144. https://doi.org/10.1007/s00429-019-01953-2

DeYoung, C. G. (2010). Personality neuroscience and the biology of traits. Social Personality Psychology Compass, 4(12), 1165–1180.

DeYoung, C. G., Hirsh, J. B., Shane, M. S., et al. (2010). Testing predictions from personality neuroscience. Brain structure and the big five. Psychol Sci, 21(6), 820–828. https://doi.org/10.1177/0956797610370159

Elliott, M. L., Knodt, A. R., Cooke, M., et al. (2019). General functional connectivity: Shared features of resting-state and task fMRI drive reliable and heritable individual differences in functional brain networks. Neuroimage, 189, 516–532. https://doi.org/10.1016/j.neuroimage.2019.01.068

Fransson, P., & Marrelec, G. (2008). The precuneus/posterior cingulate cortex plays a pivotal role in the default mode network: Evidence from a partial correlation network analysis. Neuroimage, 42(3), 1178–1184. https://doi.org/10.1016/j.neuroimage.2008.05.059

Freitas-Ferrari, M. C., Hallak, J. E., Trzesniak, C., et al. (2010). Neuroimaging in social anxiety disorder: a systematic review of the literature. Prog Neuropsychopharmacol Biol Psychiatry, 34(4), 565–580. https://doi.org/10.1016/j.pnpbp.2010.02.028

Frith, C. D. (2007). The social brain? Philos Trans R Soc Lond B Biol Sci, 362(1480), 671–678. https://doi.org/10.1098/rstb.2006.2003

Gaviria, J., Rey, G., Bolton, T., et al. (2021). Dynamic functional brain networks underlying the temporal inertia of negative emotions. Neuroimage, 240, 118377. https://doi.org/10.1016/j.neuroimage.2021.118377

Gilmore, A. W., Nelson, S. M., & McDermott, K. B. (2015). A parietal memory network revealed by multiple MRI methods. Trends Cogn Sci, 19(9), 534–543. https://doi.org/10.1016/j.tics.2015.07.004

Hecht, E. E., Robins, D. L., Gautam, P., et al. (2017). Intranasal oxytocin reduces social perception in women: Neural activation and individual variation. Neuroimage, 147, 314–329. https://doi.org/10.1016/j.neuroimage.2016.12.046

Horta, M., Ziaei, M., Lin, T., et al. (2019). Oxytocin alters patterns of brain activity and amygdalar connectivity by age during dynamic facial emotion identification. Neurobiol Aging, 78, 42–51. https://doi.org/10.1016/j.neurobiolaging.2019.01.016

Hurlemann, R., Patin, A., Onur, O. A., et al. (2010). Oxytocin enhances amygdala-dependent, socially reinforced learning and emotional empathy in humans. J Neurosci, 30(14), 4999–5007. https://doi.org/10.1523/JNEUROSCI.5538-09.2010

Hyon, R., Youm, Y., Kim, J., et al. (2020). Similarity in functional brain connectivity at rest predicts interpersonal closeness in the social network of an entire village. Proceedings of the National Academy of Sciences of the United States of America, 117(52), 33149–33160. https://doi.org/10.1073/pnas.2013606117

Jiang, X., Ma, X. L., Geng, Y. Y., et al. (2021). Intrinsic, dynamic and effective connectivity among large-scale brain networks modulated by oxytocin. Neuroimage, 227, 117668. https://doi.org/10.1016/j.neuroimage.2020.117668

Kircher, T. T., Senior, C., Phillips, M. L., et al. (2000). Towards a functional neuroanatomy of self processing: effects of faces and words. Brain Res Cogn Brain Res, 10(1-2), 133–144. https://doi.org/10.1016/s0926-6410(00)00036-7

Kjaer, T. W., Nowak, M., & Lou, H. C. (2002). Reflective self-awareness and conscious states: PET evidence for a common midline parietofrontal core. Neuroimage, 17(2), 1080–1086. https://www.ncbi.nlm.nih.gov/pubmed/12377180

Koch, S. B., van Zuiden, M., Nawijn, L., et al. (2016). Intranasal Oxytocin Normalizes Amygdala Functional Connectivity in Posttraumatic Stress Disorder. Neuropsychopharmacology, 41(8), 2041–2051. https://doi.org/10.1038/npp.2016.1

Kong, R., Li, J., Orban, C., et al. (2019). Spatial Topography of Individual-Specific Cortical Networks Predicts Human Cognition, Personality, and Emotion. Cerebral Cortex, 29(6), 2533–2551. https://doi.org/10.1093/cercor/bhy123

Kumsta, R., & Heinrichs, M. (2013). Oxytocin, stress and social behavior: neurogenetics of the human oxytocin system. Curr Opin Neurobiol, 23(1), 11–16. https://doi.org/10.1016/j.conb.2012.09.004

Laumann, T. O., Gordon, E. M., Adeyemo, B., et al. (2015). Functional System and Areal Organization of a Highly Sampled Individual Human Brain. Neuron, 87(3), 657–670. https://doi.org/10.1016/j.neuron.2015.06.037

Lee, S., Parthasarathi, T., & Kable, J. W. (2021). The Ventral and Dorsal Default Mode Networks Are Dissociably Modulated by the Vividness and Valence of Imagined Events. J Neurosci, 41(24), 5243–5250. https://doi.org/10.1523/JNEUROSCI.1273-20.2021

Lopatina, O. L., Komleva, Y. K., Gorina, Y. V., et al. (2018). Neurobiological Aspects of Face Recognition: The Role of Oxytocin. Front Behav Neurosci, 12, 195. https://doi.org/10.3389/fnbeh.2018.00195

Lou, H. C., Luber, B., Crupain, M., et al. (2004). Parietal cortex and representation of the mental Self. Proc Natl Acad Sci U S A, 101(17), 6827–6832. https://doi.org/10.1073/pnas.0400049101

Maier, A., Scheele, D., Spengler, F. B., et al. (2019). Oxytocin reduces a chemosensory-induced stress bias in social perception. Neuropsychopharmacology, 44(2), 281–288. https://doi.org/10.1038/s41386-018-0063-3

Markett, S., Montag, C., & Reuter, M. (2018). Network Neuroscience and Personality. Personal Neurosci, 1, e14. https://doi.org/10.1017/pen.2018.12

Mennes, M., Kelly, C., Zuo, X. N., et al. (2010). Inter-individual differences in resting-state functional connectivity predict task-induced BOLD activity. Neuroimage, 50(4), 1690–1701. https://doi.org/10.1016/j.neuroimage.2010.01.002

Meyer, M. C., van Oort, E. S., & Barth, M. (2013). Electrophysiological correlation patterns of resting state networks in single subjects: a combined EEG–fMRI study. Brain topography, 26(1), 98–109.

Min, J., & Wang, J. (2017). Analysis of Sleep Signals Based on Permutation Symbolic Transfer Entropy. Proceedings of the 3rd Annual International Conference on Electronics, Electrical Engineering and Information Science (Eeeis 2017), 131, 365–371.

Pang, L., Li, H., Liu, Q., et al. (2022). Resting-state functional connectivity of social brain regions predicts motivated dishonesty. Neuroimage, 256, 119253. https://doi.org/10.1016/j.neuroimage.2022.119253

Parkinson, C., Kleinbaum, A. M., & Wheatley, T. (2018). Similar neural responses predict friendship. Nat Commun, 9(1), 332. https://doi.org/10.1038/s41467-017-02722-7

Pezzulo, G., Zorzi, M., & Corbetta, M. (2021). The secret life of predictive brains: what’s spontaneous activity for? Trends in Cognitive Sciences, 25(9), 730–743.

Rachakonda, S., Egolf, E., Correa, N., et al. (2007). Group ICA of fMRI toolbox (GIFT) manual. Dostupnez [cit 2011-11-5].

Reineberg, A. E., Andrews-Hanna, J. R., Depue, B. E., et al. (2015). Resting-state networks predict individual differences in common and specific aspects of executive function. Neuroimage, 104, 69–78. https://doi.org/10.1016/j.neuroimage.2014.09.045

Scheele, D., Wille, A., Kendrick, K. M., et al. (2013). Oxytocin enhances brain reward system responses in men viewing the face of their female partner. Proceedings of the National Academy of Sciences of the United States of America, 110(50), 20308–20313. https://doi.org/10.1073/pnas.1314190110

Shine, J. M., Breakspear, M., Bell, P. T., et al. (2019). Human cognition involves the dynamic integration of neural activity and neuromodulatory systems. Nature neuroscience,22(2), 289–296.

Shirer, W. R., Ryali, S., Rykhlevskaia, E., et al. (2012). Decoding subject-driven cognitive states with whole-brain connectivity patterns. Cerebral Cortex, 22(1), 158–165. https://doi.org/10.1093/cercor/bhr099

Simony, E., Honey, C. J., Chen, J., et al. (2016). Dynamic reconfiguration of the default mode network during narrative comprehension. Nat Commun, 7(1), 12141. https://doi.org/10.1038/ncomms12141

Stanley, D. A., & Adolphs, R. (2013). Toward a neural basis for social behavior. Neuron, 80(3), 816–826. https://doi.org/10.1016/j.neuron.2013.10.038

Tavor, I., Parker Jones, O., Mars, R. B., et al. (2016). Task-free MRI predicts individual differences in brain activity during task performance. Science, 352(6282), 216–220. https://doi.org/10.1126/science.aad8127

Thompson, E. R. (2007). Development and validation of an internationally reliable short-form of the positive and negative affect schedule (Panas). Journal of Cross-Cultural Psychology, 38(2), 227–242. https://doi.org/10.1177/0022022106297301

Utevsky, A. V., Smith, D. V., & Huettel, S. A. (2014). Precuneus Is a Functional Core of the Default-Mode Network. Journal of Neuroscience, 34(3), 932–940. https://doi.org/10.1523/Jneurosci.4227-13.2014

Vogeley, K., Bussfeld, P., Newen, A., et al. (2001). Mind reading: neural mechanisms of theory of mind and self-perspective. Neuroimage, 14(1 Pt 1), 170–181. https://doi.org/10.1006/nimg.2001.0789

Vogeley, K., & Fink, G. R. (2003). Neural correlates of the first-person-perspective. Trends Cogn Sci, 7(1), 38–42. https://doi.org/10.1016/s1364-6613(02)00003-7

Wang, Y., Wang, R., & Wu, H. (2022). The role of oxytocin in modulating self–other distinction in human brain: a pharmacological fMRI study. Cerebral Cortex.

Wu, H., Feng, C., Lu, X., et al. (2020). Oxytocin effects on the resting-state mentalizing brain network. Brain Imaging Behav, 14(6), 2530–2541. https://doi.org/10.1007/s11682-019-00205-5

Wu, H., Liu, X., Hagan, C. C., et al. (2020). Mentalizing during social InterAction: A four component model. Cortex, 126, 242–252. https://doi.org/10.1016/j.cortex.2019.12.031

Xin, F., Zhou, F., Zhou, X. Q., et al. (2021). Oxytocin Modulates the Intrinsic Dynamics Between Attention-Related Large-Scale Networks. Cerebral Cortex, 31(3), 1848–1860. https://doi.org/10.1093/cercor/bhy295

Xin, F., Zhou, X., Dong, D., et al. (2020). Oxytocin Differentially Modulates Amygdala Responses during Top-Down and Bottom-Up Aversive Anticipation. Adv Sci (Weinh), 7(16), 2001077. https://doi.org/10.1002/advs.202001077

Zadbood, A., Chen, J., Leong, Y. C., et al. (2017). How We Transmit Memories to Other Brains: Constructing Shared Neural Representations Via Communication. Cerebral Cortex, 27(10), 4988–5000. https://doi.org/10.1093/cercor/bhx202

Zheng, S., Liang, Z., Qu, Y., et al. (2022). Kuramoto Model-Based Analysis Reveals Oxytocin Effects on Brain Network Dynamics. Int J Neural Syst, 32(2), 2250002. https://doi.org/10.1142/S0129065722500022

Zheng, S., Punia, D., Wu, H., et al. (2021). Graph Theoretic Analysis Reveals Intranasal Oxytocin Induced Network Changes Over Frontal Regions. Neuroscience, 459, 153–165. https://doi.org/10.1016/j.neuroscience.2021.01.018

Zhu, R., Liu, C., Li, T., et al. (2019). Intranasal oxytocin reduces reactive aggression in men but not in women: A computational approach. Psychoneuroendocrinology, 108, 172–181.

