## supplementary materials for "Oxytocin modulates social brain network correlations in resting and task state"

**Supplementary file**

Table S1: Predictive accuracy of others using two methods.

| **SM OM t-value** | | | | **p** |
| --- | --- | --- | --- | --- |
| PL DDMN | 0.35 | 0.33 | 2.28 | 0.026 |
| PL PN | 0.37 | 0.32 | 4.44 | 0.000 |
| PL VDMN | 0.34 | 0.32 | 1.38 | 0.174 |
| PL SN | 0.24 | 0.22 | 2.22 | 0.031 |
| OT ASN | 0.22 | 0.21 | 1.96 | 0.054 |
| OT PN | 0.40 | 0.36 | 3.42 | 0.001 |
| OT VDMN | 0.34 | 0.33 | 2.60 | 0.012 |
| OT SN | 0.24 | 0.22 | 3.33 | 0.002 |

*Note:* Means(standard deviations), and two-tailed t-test (direction: SM - OM) results of the comparisons on Predictive accuracy.

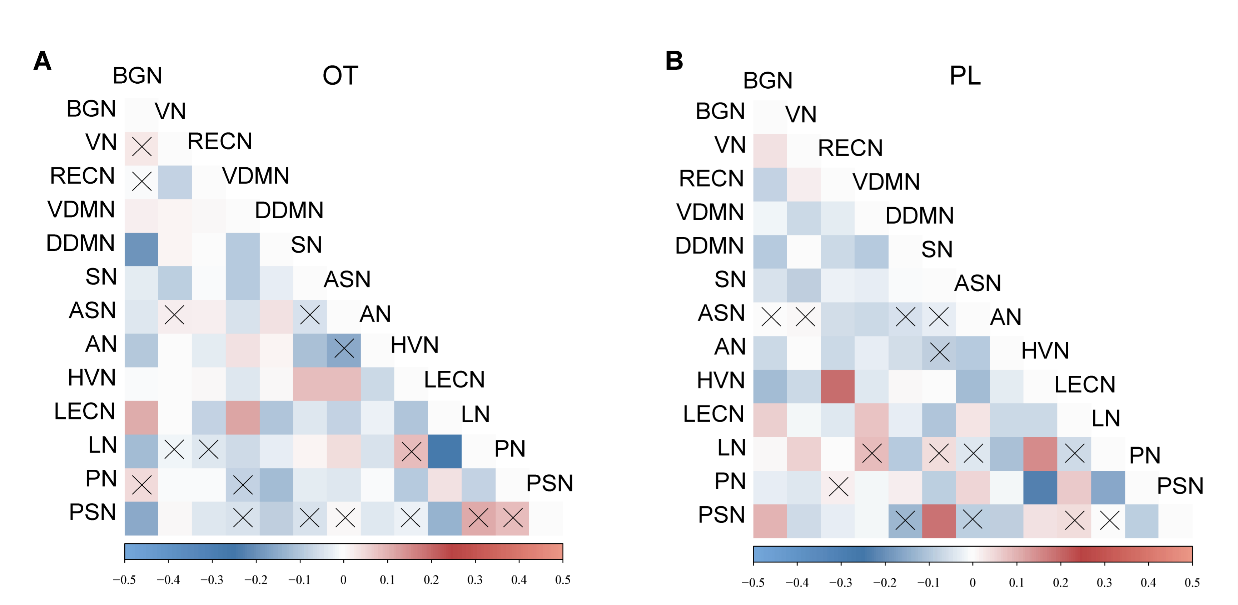

Figure S1 Correlation matrix between brain networks. A) Correlation matrix for the OT group. B) Correlation matrix for the PL group. The black marks mean the correlation coefficients are not significant($P_{FDR}>0.05$).

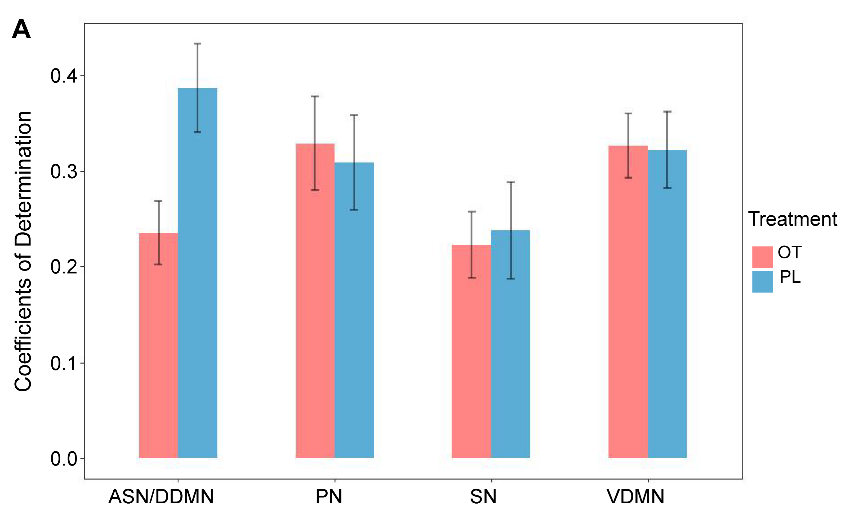

Figure S2 The models’ coefficients of determination($R^{2}$). We averaged the coefficients of all subjects for each network. In the bar with ASN/DDMN horizontal coordinate, the red bar represents the mean coefficients of ASN, and the blue bar represents the mean coefficients of DDMN.

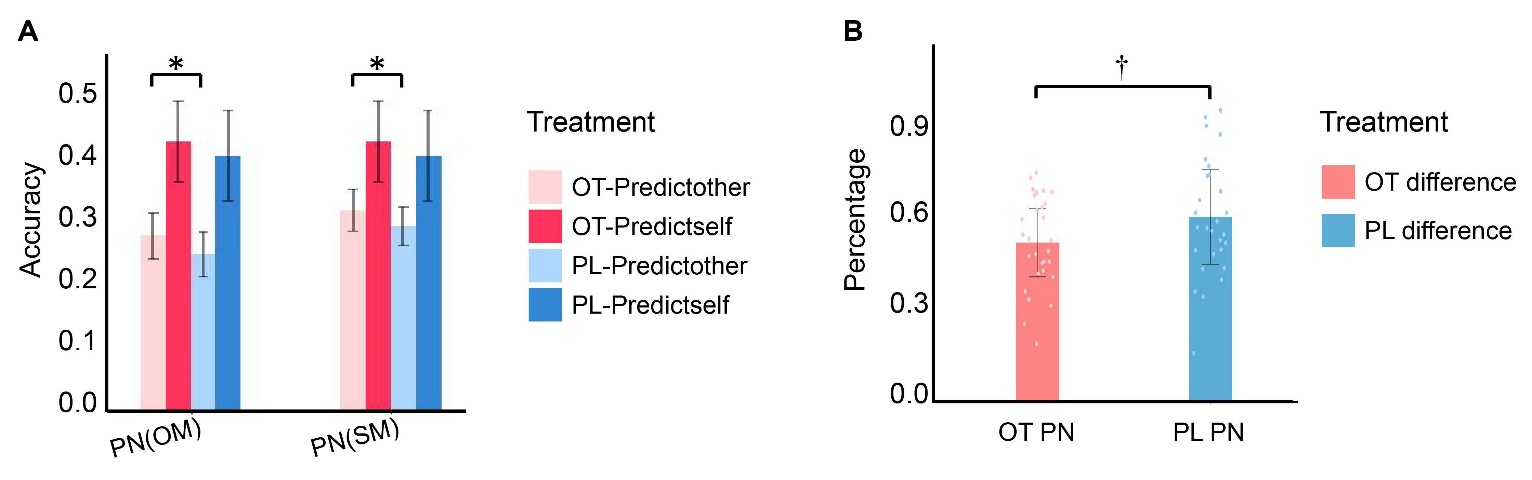

Figure S3 Performance of the model and specificity of the individual predictions. A) This figure shows the accuracy of using one’s own model (SM) and others’ models (OM) to predict one’s own (deep color) and others’ task-evoked network activation (light color) for both conditions. $t_{OM}\left( 58 \right)=2.29, p=0.03;t_{SM}\left( 58 \right)=2.15, p=0.036.$ B) The bar plot, which was produced by the method using OM, indicates the difference between the accuracy of using participants’ resting-state data to predict their task activation and predicting others’ activation (as a percentage relative to the average accuracy of predicting others). $t\left( 58 \right)=1.77, p=0.08.The error bars show 1.96 standard deviations.$

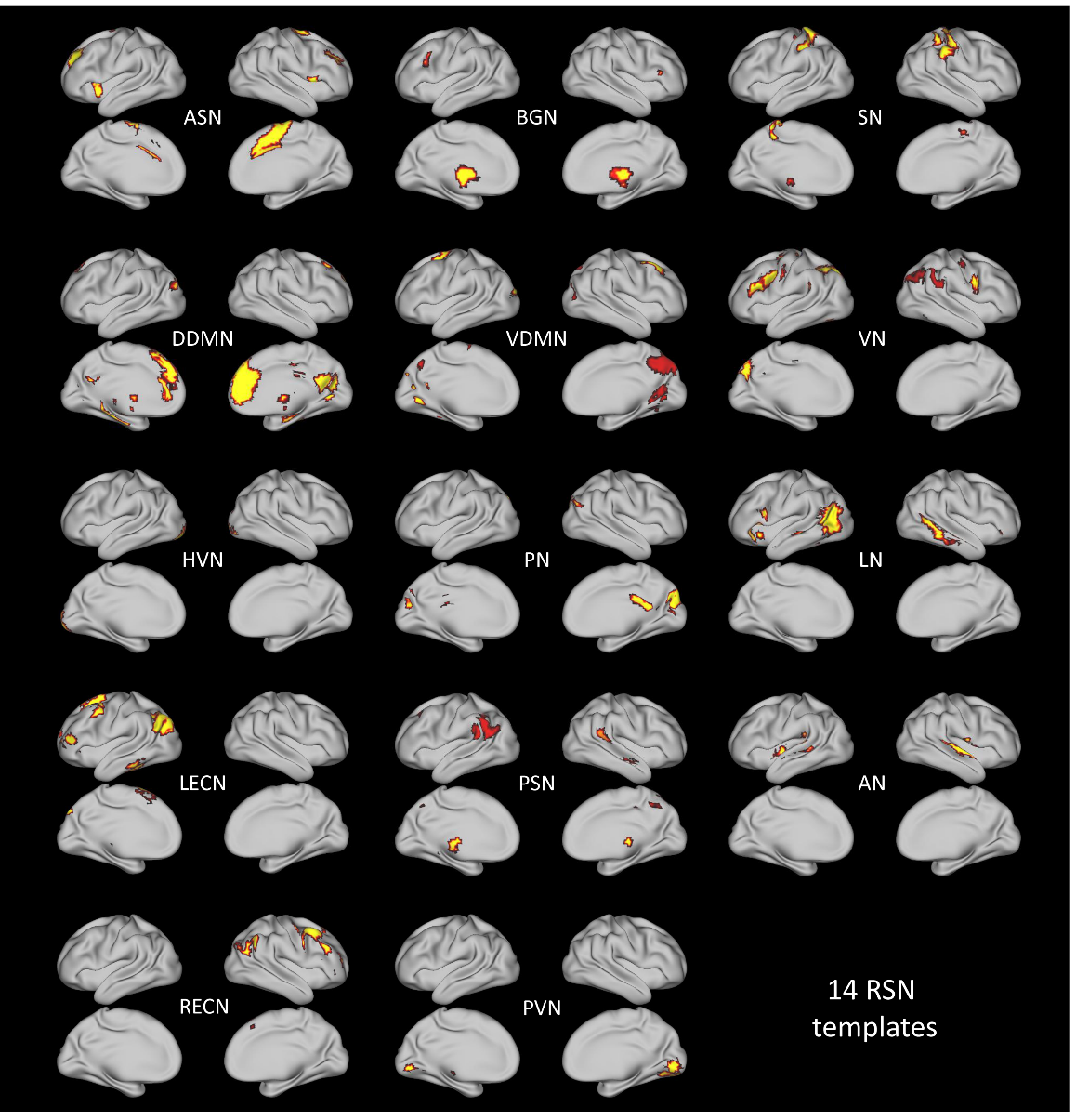

Figure S4 The regions contained with each RSN template. ASN: Insula/Dorsal Anterior Cingulate Cortex (dACC), BGN: Basal Ganglia, SN: Sensorimotor, DDMN: Posterior Cingulate Cortex (PCC)/Medial Prefrontal Cortex (MPFC), VDMN: Retrosplenial Cortex (RSC)/Medial Temporal Lobe (MTL), VN: Intraparietal Sulcus (IPS)/Frontal Eye Field (FEF), HVN: Secondary Visual Cortex (V2), PN: Precuneus, LN: Language, LECN: Left Dorsolateral Prefrontal Cortex (DLPFC)/Left Parietal Lobe, PSN: Posterior Insula, AN: Auditory, RECN: Right Dorsolateral Prefrontal Cortex (DLPFC)/Right Parietal Lobe, PVN: Primary Visual Cortex (V1).
